# On the recovery of malformed horseshoe crabs across multiple moulting stages

**DOI:** 10.1101/2024.03.27.587089

**Authors:** Russell D. C. Bicknell, Carmela Cuomo

## Abstract

Malformed horseshoe crabs have been documented for over a century. However, most of these records are anecdotal observations of often striking morphologies recorded in isolation. There is therefore little understanding of how malformations are manifested and how they can develop in the group. Here we consider the moult sequences of three extant *Limulus polyphemus* individuals to explore different patterns of malformation development. One specimen with an injured telson demonstrates a gradual recovery of the telson section over three moulting events. The second individual demonstrates a fused thoracetron-telson articulation with a hole for the telson. This individual shows consistent growth of a reduced telson across moults. The third individual shows a thoracetronic injury incurred during at least moult-stage 7 that shows no evidence of recovery over five moulting stages. These records illustrate that horseshoe crab malformation recovery is far more complicated than previously thought. This also suggests that unless an exoskeletal section has functional morphological importance (i.e., the telson), the region is unlikely to recover from an older malformation. From a conservation standpoint, the ability or inability to fully recover from injury affects a horseshoe crab’s ability to survive and/or reproduce in the wild particularly if the injury affects the telson. Given the global decline in horseshoe crab populations and conservation efforts underway, the extent of injuries in extant populations of horseshoe crabs may affect population recovery and should be considered.

## Introduction

Horseshoe crab represent rare examples of modern marine chelicerates (Størmer 1955; Sekiguchi 1988; Dunlop and Lamsdell 2017). This unique phylogenetic position, their use in the biomedical industry (Gokudan et al. 1999; Shuster Jr. et al. 2003; Akbar John et al. 2011; Krisfalusi-Gannon et al. 2018), and their ecological importance (Sokoloff 1978; Shuster Jr. 1982; Botton and Ropes 1987, 1989; Sekiguchi 1988; Patil and Anil 2000; Pierce et al. 2000; Shuster Jr. 2001; Sekiguchi and Shuster Jr. 2009; Shuster Jr. and Sekiguchi 2009) have driven research into modern species. This interest is compounded by the exceptionally long evolutionary history of Xiphosura (Størmer 1952; Eldredge 1974; Anderson and Shuster Jr. 2003; Rudkin et al. 2008; Lamsdell 2016; Bicknell et al. 2022a), the xiphosuran origin deep in the Paleozoic (Rudkin et al. 2008; Rudkin and Young 2009; Bicknell and Pates 2020), the two major radiations (Lamsdell 2016; Bicknell et al. 2021a; Bicknell and Shcherbakov 2021; Lustri et al. 2021), and the evidence of conserved dorsal (Barthel 1974; Fisher 1984; Avise et al. 1994; Eldredge et al. 2005), ventral (Bicknell et al. 2019b) and internal (Briggs et al. 2005; Bicknell et al. 2021c, 2022d) morphologies. The morphological similarity to extinct arthropods, such as trilobites and sea scorpions, has resulted in horseshoe crabs being applied to understanding the paleobiology of fossil forms (Brandt 2002; Paterson et al. 2008; Bicknell et al. 2018a. 2019a, 2021b, 2022c). In particular, horseshoe crabs have been used to understand malformation recovery in trilobites (Bicknell et al. 2018b, 2022b Bicknell and Pates 2019). However, beyond one anecdotal example (Bicknell et al. 2022b), there is effectively no record of xiphosuran recovery from malformations. This likely reflects the slow rate of horseshoe crab moulting and the high likelihood of difficulty moulting, possibly ending in death (Vosatka 1970; Fisher 1977; Sokoloff 1978; Botton and Loveland 1989; Penn and Brockmann 1995). Injury to the telson may also make the animal less able to right itself when flipped, making it vulnerable to predation and death. As a result, the appropriateness of this analogue is uncertain. To explore patterns of malformation recovery, moult sequences of three *Limulus polyphemus* individuals are considered. These specimens record how thoracetronic and telson malformations change across multiple moulting events.

## Methods

Moults of three horseshoe crabs collected by C.C. were considered. The first specimen, with injuries to its thoracetron and lacking a telson was found during a collection trip and brought to the lab by students. The other two specimens were reared in the lab from eggs legally collected from New Haven Harbor, New Haven, Connecticut. All specimens were reared in the laboratory according to the proprietary protocols of Limulus Ranch, LLC. Water parameters, including salinity, ammonia, phosphate, calcium, alkalinity, and dissolved oxygen were checked weekly. Animals were fed ad libitum. Moulting events were noted, and molts were removed from the tanks and allowed to air dry prior to being stored. The wild caught injured horseshoe crab molted during its second year in the lab and then again in its third year; it died during its fourth year in captivity. One of the two laboratory hatched and reared animals also died after several years and numerous successful molts. The dead animals were preserved in 10% formalin and then transferred to 70% ethanol for storage. The moults and carcasses of these two individuals were catalogued into the Yale Peabody Museum Invertebrate Zoological (YPM IZ) collection. The third animal, hatched in 2020, is still alive; its moults were also catalogued into the YPM IZ collections. The moults, carcasses, and the live specimen were photographed under LED lights with an Olympus E-M1MarkIII camera with a 12–45 mm lens. Images were stacked using OM Capture. Measurements of specimens were gathered using digital calipers (Tables 1, 2). Proposed growth stages for specimens follow Lamsdell (2020).

**Table 1:** Measurements of growth series of *Limulus polyphemus* showing malformed telsons. Percentage of telson growth is presented.– indicates that the % value could not be calculated. — indicates the specimen was not catalogued, as it is still alive.

| Specimen | Growth stage | Prosomal length (mm) | Prosomal width (mm) | Thoracetrone length (mm) | Thoracetrone width (mm) | Telson length (mm) | % telson recovery | Figure |
| --- | --- | --- | --- | --- | --- | --- | --- | --- |
| YPM IZ 111021 | 13 | 45.51 | 66.34 | 30.57 | 47.38 | 5.8 | — | Figure 1A–C |
| YPM IZ 111022 | 14 | 54.39 | 87.28 | 37.64 | 58.07 | 10.84 | 86.9% | Figure 1D–F |
| YPM IZ 111029 | 15 | 63.74 | 99.4 | 46.2 | 67.81 | 25.12 | 132% | Figure 1G–I |
| YPM IZ 111023 | 9 | 16.11 | 25.15 | 14.46 | 17.90 | Not observed | — | Figure 3A, B |
| YPM IZ 111024 | 10 | 20.87 | 32.91 | 19.86 | 23.21 | 7.53 | — | Figure 3C–E |
| — | 11 | 26.35 | 43.77 | 27.05 | 30.26 | 10.05 | 33.5% | Figure 3F |

**Table 2:** Measurements of growth series for *Limulus polyphemus* showing malformed thoracetron. indicates that the measurement could not be gathered due to damage to the exoskeleton or that the % value could not be calculated.

| Specimen | Growth stage | Prosomal length (mm) | Prosomal width (mm) | Thoracetron length (mm) | Thoracetron width (mm) | Telson length (mm) | Injury length (mm) | Injury growth | Figure |
| --- | --- | --- | --- | --- | --- | --- | --- | --- | --- |
| YPM IZ 111025 | 7 | 17.3 | 26.8 | 11.78 | 16.04 | 18.9 | 6.14 | – | Figure 2A, E |
| YPM IZ 111026 | 9 | 20.44 | – | 14.47 | 18.99 | 25.13 | 7.36 | 19.9% | Figure 2B, F |
| YPM IZ 111027 | 11 | 27.17 | 42.38 | 18.76 | 26.53 | 30.35 | 9.65 | 31.1% | Figure 2C, G |
| YPM IZ 111028 | 12 | 34.57 | 51.8 | 21.7 | 31.13 | 33.99 | 12.03 | 24.7% | Figure 2D, H |

## Results

The first examined individual underwent two successful moults after being brought to the lab (Table 1). The first moult, occurred nearly two years after the animal was collected and the injury presumably took place, shows marked callousing of the posterior thoracetron and near complete removal of the telson. The terminal thoracetronic spine and posterior most fixed spine on the right side are truncated into a U-shaped indentation (Figure 1A–C). The second post-injury moult shows telson recovery to 10.84 mm long; the section is deflected to the left (Figure 1D–F). The malformed terminal and posterior-most fixed spine show partial recovery and rounding. The most posterior moveable is not present. The carcass, believed to be six years of age, shows additional telson recovery to 25.12 mm and additional rounding of the malformed spines (Figure 1G–I). The moveable spine on the right side did not recover.

**FIGURE 1.**
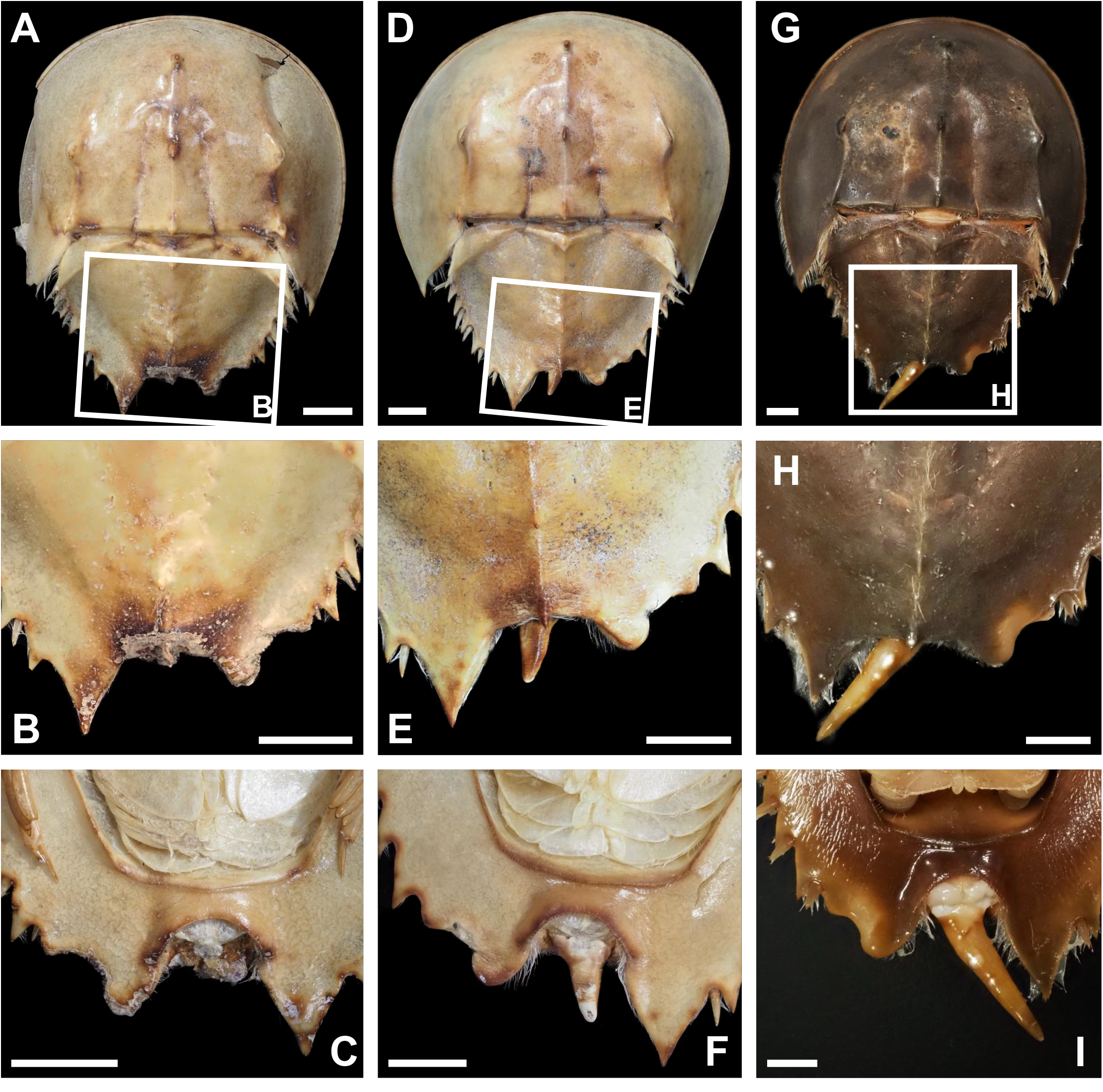
Malformed *Limulus polyphemus* showing malformed thoracetron and telson with recovery over moults. (A–C) First recorded stage, growth stage 13, moult (YPM IZ 111021). (A) Complete specimen, dorsal view. (B) Close up of box in (A). (C) Close up of malformation, ventral view. (D–F) Second recorded stage, growth stage 14, moult (YPM IZ 111022). (D) Complete specimen, dorsal view. (E) Close up of box in (D). (F) Close up of malformation, ventral view. (G–I) Third recorded stage, growth stage 15, carcass (YPM IZ111029). (G) Complete specimen, dorsal view. (H) Close up of box in (G). (I) Close up of malformation, ventral view. Scale bars: 10 mm

The second individual shows a thoracetronic malformation over multiple moults (Figure 2, Table 2). A W-shaped indentation on the right lateral lobe is observed where moveable and fixed spines 1–4 are located (Shuster Jr. 1982). While the malformation is more pronounced in subsequent moult-stages (compare Figure 2A, to Figure 2D), the malformation border has a comparable thickness to other regions along the thoracetron border. The malformation increases in length between 6.14–12.03 mm across moulting events, indicating an increase in size scaled with growth after moulting. No evidence for malformation recovery, such as development of movable or fixed spines, is observed across moults.

**FIGURE 2.**
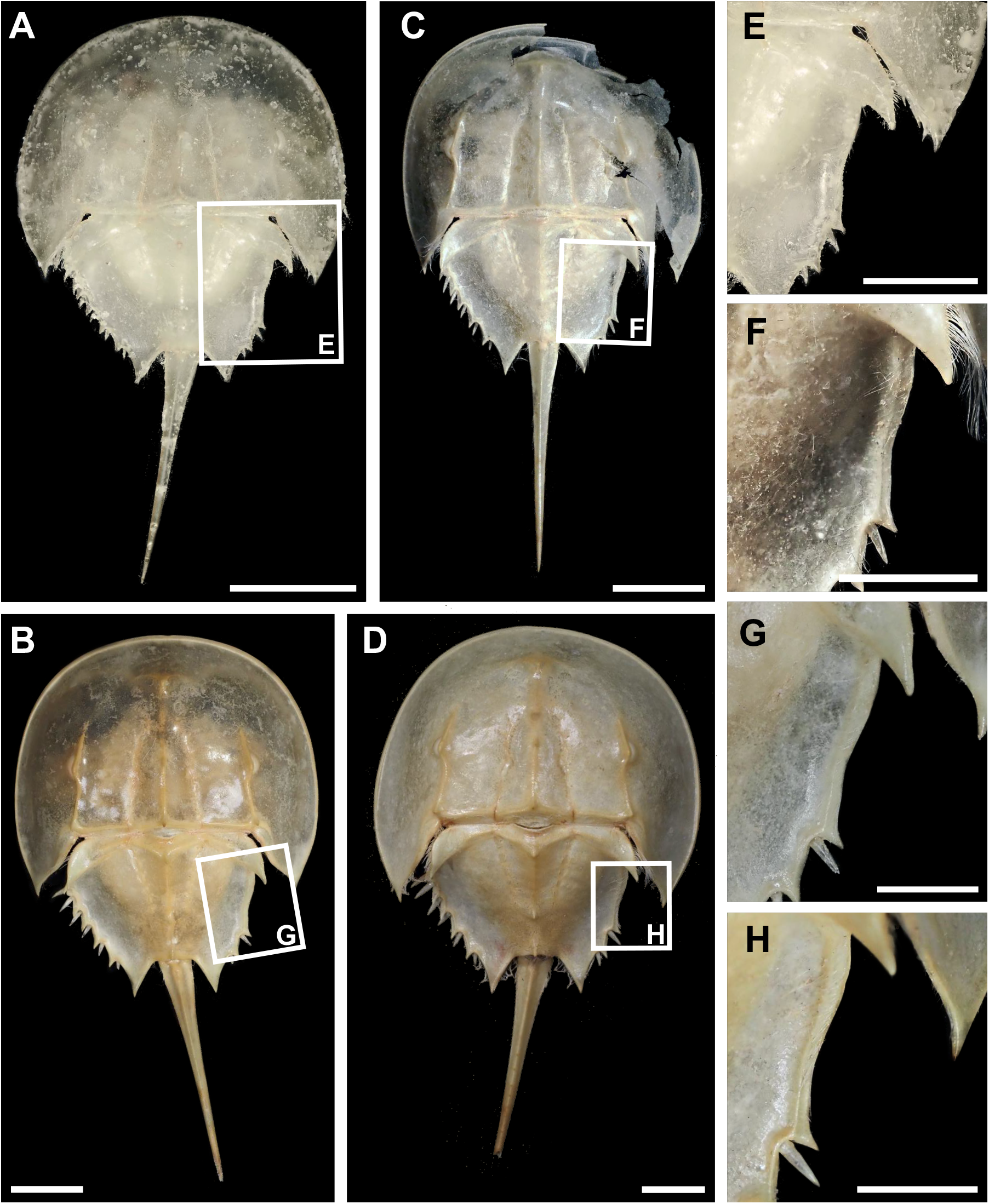
Malformed *Limulus polyphemus* moults showing W-shaped indentation in thoracetron that shows no recovery. (A, E) First recorded stage, growth stage 7 (YPM IZ 111025). (A) Complete specimen. (E) Close up of box in (A). (B, F) Second recorded stage, growth stage 9, (YPM IZ 111026). (B) Complete specimen. (F) Close up of box in (B). (C, G) Second recorded stage, growth stage 11, (YPM IZ 111027). (C) Complete specimen. (G) Close up of box in (C). (D, H) Third recorded stage, growth stage 12, (YPM IZ 111028). (D) Complete specimen. (H) Close up of box in (D). Scale bars: (A–D): 10 mm; (E–H): 5 mm.

The third individual, now three years of age, shows fusion of the telson embayment (Figure 3). In the moults and the current growth stage, the thoracetron lacks a telson articulation. In the first preserved moult (Stage 9), a hole in the dorsal thoracetron is observed, but no telson noted. The anus is located approximately where the telson embayment would be expected. The next preserved moult (Stage 10) shows the development of a thin, 7.5 mm long telson from this hole. In its current stage (Stage 11) the individual has 10.05 mm long telson, indicating limited growth associated with moulting.

**FIGURE 3.**
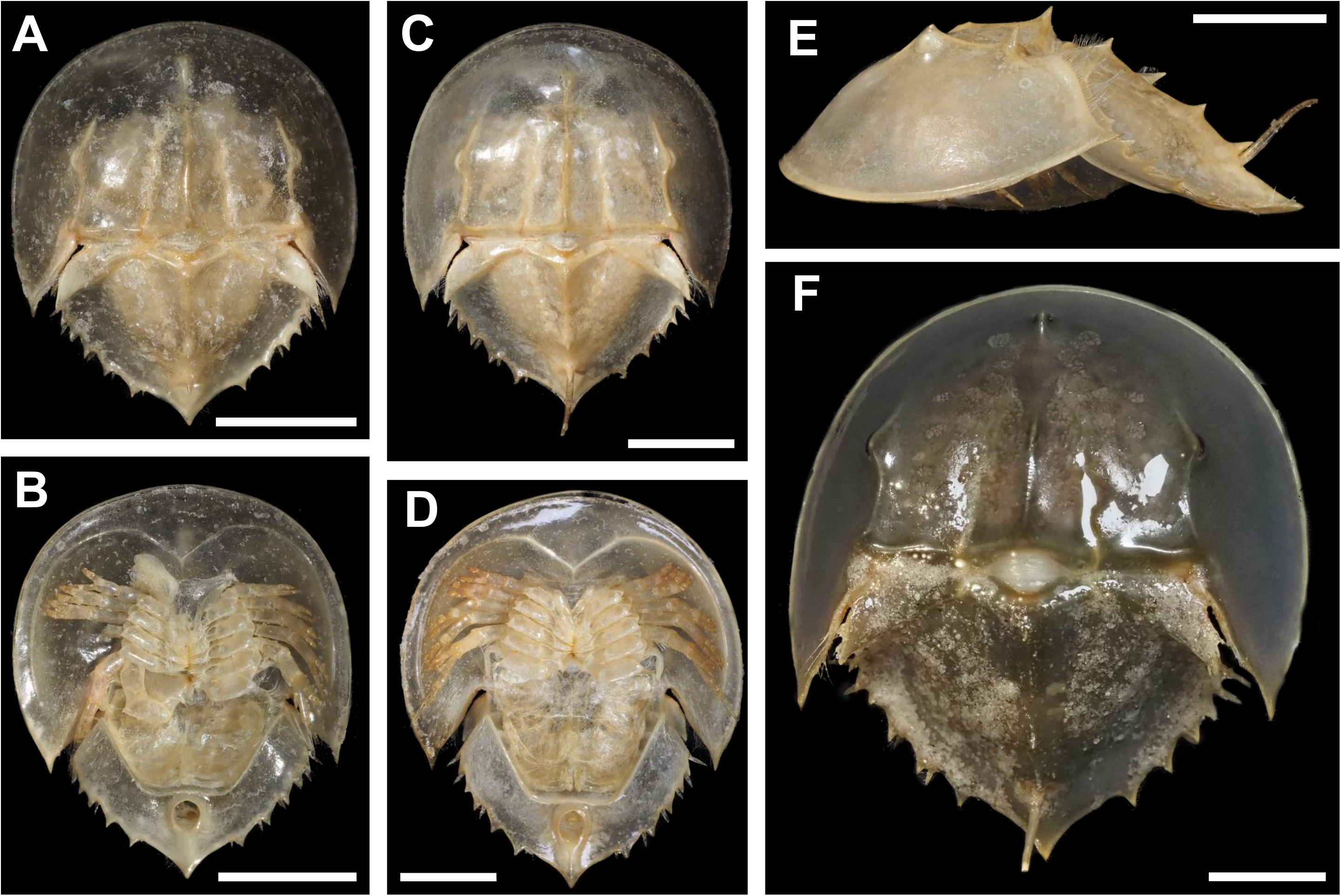
Malformed *Limulus polyphemus* showing fusion of the thoracetron and development of reduced telson. (A, B) First recorded stage, growth stage 9, moult (YPM IZ 111023). (A) Dorsal view. (B) Ventral view. (C–E) Second recorded stage, growth stage 10, moult (YPM IZ 111024). (C) Dorsal view. (D) Ventral view. (E) Lateral view showing deformed telson. (F) Third recorded stage, growth stage 11, alive specimen, no catalogued. Scale bars: All 10 mm.

## Discussion

Malformations in arthropods are commonly classed into three main divisions: injuries, teratologies, and infestation (Owen 1985; Babcock 1993, 2003; Bicknell and Paterson 2018; Bicknell et al. 2018b; Bicknell and Smith 2022; De Baets et al. 2022). Injuries reflect exoskeletal break or damage that is repaired over moult events. These can show evidence of recovery (Rudkin 1979, 1985; Babcock 1993). Teratologies record abnormal developments associated with genetic and development complications, often resulting in bizarre, abnormal morphologies (Owen 1985; Pandourski and Evtimova 2005; Pandourski and Evtimova 2009; Bicknell and Smith 2021; Zong 2021). Finally, infestations are often observed as rounded structures in the exoskeleton, indicative of pathological neoplasms (Šnajdr 1978; De Baets et al. 2022). Within the framework, we present examples of injuries (Figures 2) and teratological development (Figure 3).

Two injuries are recorded here–the injured and recovered telson and posterior thoracetron in YPM IZ 111021, 111022, 111029 and the W-shaped indentation in YPM IZ 111025–111028. The injured telson in YPM IZ 111021, 111022, 111029 shows 89–132% of the telson length growth compared to previous moult stages. This illustrates a substantial, albeit slow, telson recovery. Over more moulting events, the telson length may have fully recovered. This example contrasts the more marked recovery of an injured telson observed in a wild *Tachypleus tridentatus* that is thought to possibly reflect one moult event (Bicknell et al. 2022b). The telson recovery process may be species dependent or reflect the raising of the individual in a controlled environment.

In considering the thoracetronic injury, we demonstrate the first record of an injury that did not recover over moults–the moveable and fixed spines did not show any regeneration. This contrasts the pattern observed in the horseshoe crab telson and demonstrates that functionally less important exoskeletal regions may not have been assigned sufficient resources allotted for recovery. Further, injury size increases with each moult stage, demonstrating that thoracetronic injuries can scale with animal size without showing regeneration. As such, other horseshoe crab thoracetron injuries (see Jell 1989; Bicknell et al. 2018b, 2022b; Bicknell and Pates 2019; Das et al. 2021) could reflect events incurred earlier in the life cycle that did not recover.

The documented teratology consists of the thoracetron-telson articulation fusion in YPM IZ 10023, 10024. Due to this developmental malfunction, the individual developed a hole on the dorsal thoracetron for the reduced telson. The observed moult stages for this individual demonstrate that, while there is an increased telson length over moulting events, the telson width and length is much reduced compared to standard specimens (see Shuster Jr. 1982; Shuster Jr. et al. 2003; Shuster Jr. and Sekiguchi 2003). This reflects the minute diameter of the telson hole. The reduced size has presented complications for the individual, as the individual has been observed overturned and unable to right itself with the reduced telson (CC pers. Obs.). In a wild population this individual would likely not survive.

Horseshoe crab telsons are arguably the most striking morphology of the animals. As such, telson malformations have a long historical record of documentation (Packard 1870; Bateson 1894; Osburn 1911; Gravier 1930; Gudger 1935; van der Meer Mohr 1941; Shuster Jr. 1982; Bicknell et al. 2018b) and bifurcated telsons in adults are well-documented (Osburn 1911; Gudger 1935; Shuster Jr. 1982; Bicknell et al. 2018b). These morphologies have been considered injuries or genetic malformations (Gudger 1935). Although we did not observe these malformations, we propose that bifurcated telsons in adults likely reflect injuries that were incurred proximal to the thoracetron that then recover abnormally. Contrastingly, anecdotal observations by C.C. include rare examples of stage 1 and 2 moults with bifurcated telsons. In these cases, the early development of the specimens raised in laboratory conditions excludes injuries as explanations, indicating genetic malfunctions can cause these morphologies.

The application of horseshoe crab injuries to model possible recovery in trilobites has been proposed (Bicknell et al. 2018b; Bicknell and Pates 2019). This reflected the overall morphological similarity, comparable nature of injury shapes, and proposed similarity in recovery patterns. Examination of the moult series here demonstrates that the injuries along the thoracetron can lack recovery, indicating that horseshoe crabs are not the ideal analogue for understanding recovery in all trilobite exoskeletal sections. This difference may reflect the lack of tergite expression along the thoracetron, contrasting segment expression in trilobites. Despite this, studying injury patterns in horseshoe crab prosomal sections may uncover useful comparative data for interpreting rare injuries to trilobite cephalons (Owen 1985).

These malformations also have conservation implications. Given the inability of *Limulus* to “right” itself increases with telson injury (Botton and Loveland 1989; Penn and Brockmann 1995), these injuries, combined with the length of time for telson recovery may contribute to declines in *Limulus* populations. This is especially the case as stranding accounts for 6–10% of mortality in breeding animals (Botton and Loveland 1989; Penn and Brockmann 1995). Furthermore, while select thoracetronic injuries show no recovery across moults, these injuries may impact successful reproduction. As *Limulus* is currently listed as “Vulnerable”, any factors negatively influencing *Limulus* breeding and survival must be adopted when considering conservation studies.

## Acknowledgements

This research was funded by an MAT Program Postdoctoral Fellowship (to R.D.C.B). We would like to thank Eric Lazo-Wasem (YPM IZ) for help with collections.

